# CRISPR/Cas9-Assisted Transformation-Efficient Reaction (CRATER), a novel method for selective transformation

**DOI:** 10.1101/027664

**Authors:** L. J. Rothschild, D. T. Greenberg, J. R. Takahashi, K. A. Thompson, A. J. Maheshwari, R. E. Kent, G. McCutcheon, J. D. Shih, C. Calvet, T. D. Devlin, T. Ju, D. Kunin, E. Lieberman, T. Nguyen, F. G. Tran, D. Xiang, K. Fujishima

## Abstract

The CRISPR/Cas9 system has revolutionized genome editing by providing unprecedented DNA-targeting specificity. Here we demonstrate that this system can be applied to facilitate efficient plasmid selection for transformation as well as selective gene insertion into plasmid vectors by cleaving unwanted plasmid byproducts after restriction enzyme digestion and ligation. Using fluorescent and chromogenic proteins as reporters, we demonstrate that CRISPR/Cas9 cleavage excludes unwanted ligation byproducts and increases transformation efficiency of desired inserts from 20% up to 97% ± 3%. This CRISPR/Cas9-Assisted Transformation-Efficient Reaction (CRATER) protocol is a novel, inexpensive, and convenient method for obtaining specific cloning products.

## Introduction

Molecular cloning is a fundamental technique in molecular biology to produce plasmid constructs. Several methods currently exist to minimize or select against unwanted plasmid products created during ligation of inserts into vectors, including cross-incompatible sticky ends, X-gal blue/white screening, dephosphorylation of backbone sticky ends, the addition of antibiotics, and agarose electrophoresis/gel extraction. However, in special circumstances existing methods may be insufficient to quickly, cheaply, and effectively screen for specific cloning products. This is especially true for plasmids with compatible sticky ends. Genes of interest may include restriction sites that would otherwise be used to create incompatible sticky ends. A plasmid vector also may simply not include multiple restriction sites with incompatible sticky ends. Unwanted byproducts are also difficult to control in situations where blunt ends are used.

We developed a new method for degrading unwanted ligation products using the Cas9 nuclease from *Streptococcus pyogenes* in a one-pot reaction, enhancing transformation efficiency from 20% up to 97% ± 3%. The Cas9 protein is a component of the clustered, regularly interspaced, short palindromic repeats (CRISPR) system. The CRISPR/CRISPR-associated (Cas) system provides bacteria with acquired immunity by incorporating fragments of foreign DNA and using the transcribed CRISPR-RNA (crRNA) to guide the cleavage of matching dsDNA sequences (Bhaya, Wiedenheft). In type II CRISPR systems, a ternary complex of Cas9, crRNA, and trans-activating crRNA (tracrRNA) binds to and cleaves dsDNA sequences that match the crRNA and include a short protospacer-adjacent motif (PAM) recognized by Cas9 (Gasiunas, Jinek). In type II systems, the crRNA and tracrRNA can be combined into a single guide-RNA (sgRNA) that is sufficient to lead Cas9 to its target (Jinek). Further, the PAM sequence recognized by the *S. pyogenes* Cas9 is only three nucleotides in length (NGG), allowing this system to be easily adapted to recognize and cut a desired sequence (Mojica).

With the knowledge that Cas9 can be used to cleave short (~24 bp) sequences, we investigated whether this system could be adapted to cleave unwanted ligation byproducts. We used the RFP BioBrick plasmid BBa_J04450 (Supplementary fTable S1) as a starting vector and replaced the RFP insert with various genes of interest using restriction enzyme digestion and ligation, before transforming into *Escherichia coli*. We then quantified insertion efficiency based on the presence of fluorescent and chromogenic proteins in colonies and culture. We show, for the first time to our knowledge, that Cas9 and sgRNAs can be used to increase molecular cloning efficiency by cleavage of specific, undesired ligation byproducts; we call this novel technique CRISPR/Cas9-assisted transformation-efficient reaction (CRATER).

## Results

### Cas9 cleavage of plasmids enhances transformation selectively

We first sought to verify that *in vitro* Cas9 cleavage specifically selects for and purifies a desired plasmid product from an *in vitro* mixed pool. *E. coli* is known to have dramatically lower transformation efficiency for linear DNA compared with plasmids, due to intracellular exonuclease activity (Conley). To test the ability of *in vitro* Cas9 digestion to selectively prevent the transformation of multiple plasmid vectors in a mixed pool, we prepared a mixture of four different plasmids encoding color-producing proteins: RFP, *eforRed*, *amilGFP*, and *meffBlue*. We then designed four sgRNAs to selectively target each gene and added all combinations of three sgRNAs to the mixtures of plasmids. We transformed the resulting pools into *E. Coli.*

We found that *in vitro* Cas9 digestion of plasmids significantly (p<0.001) selected against transformation into *E. coli*, both upon visual inspection (Fig. 2A) and by quantification of percentage of desired colonies on transformed plates (Fig. 2B). Cas9 cleavage was verified by gel electrophoresis (Supplementary Fig. S1). The use of multiple sgRNAs in a single Cas9 reaction did not appear to interfere with successful target cleavage.

**Figure 1.**
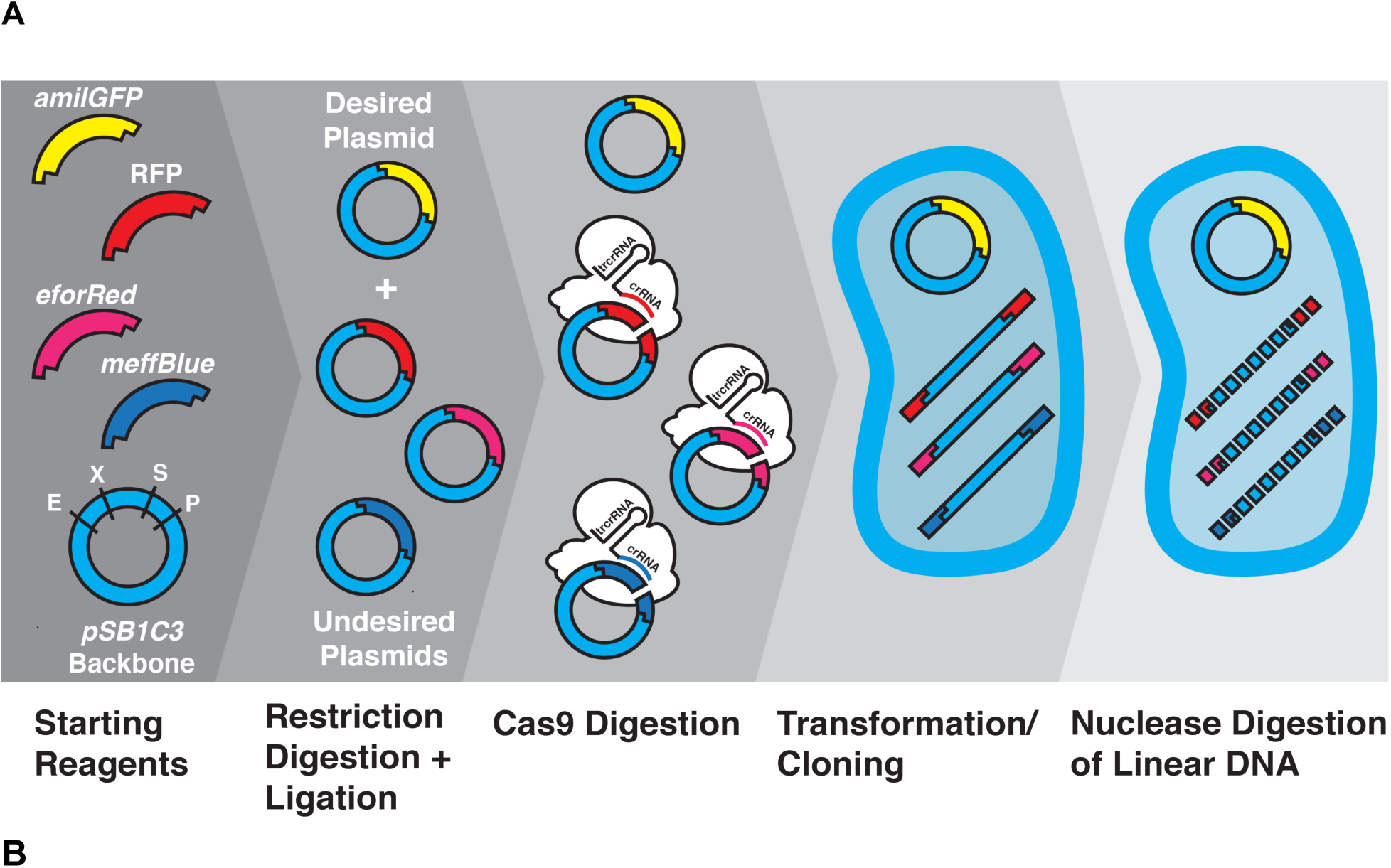
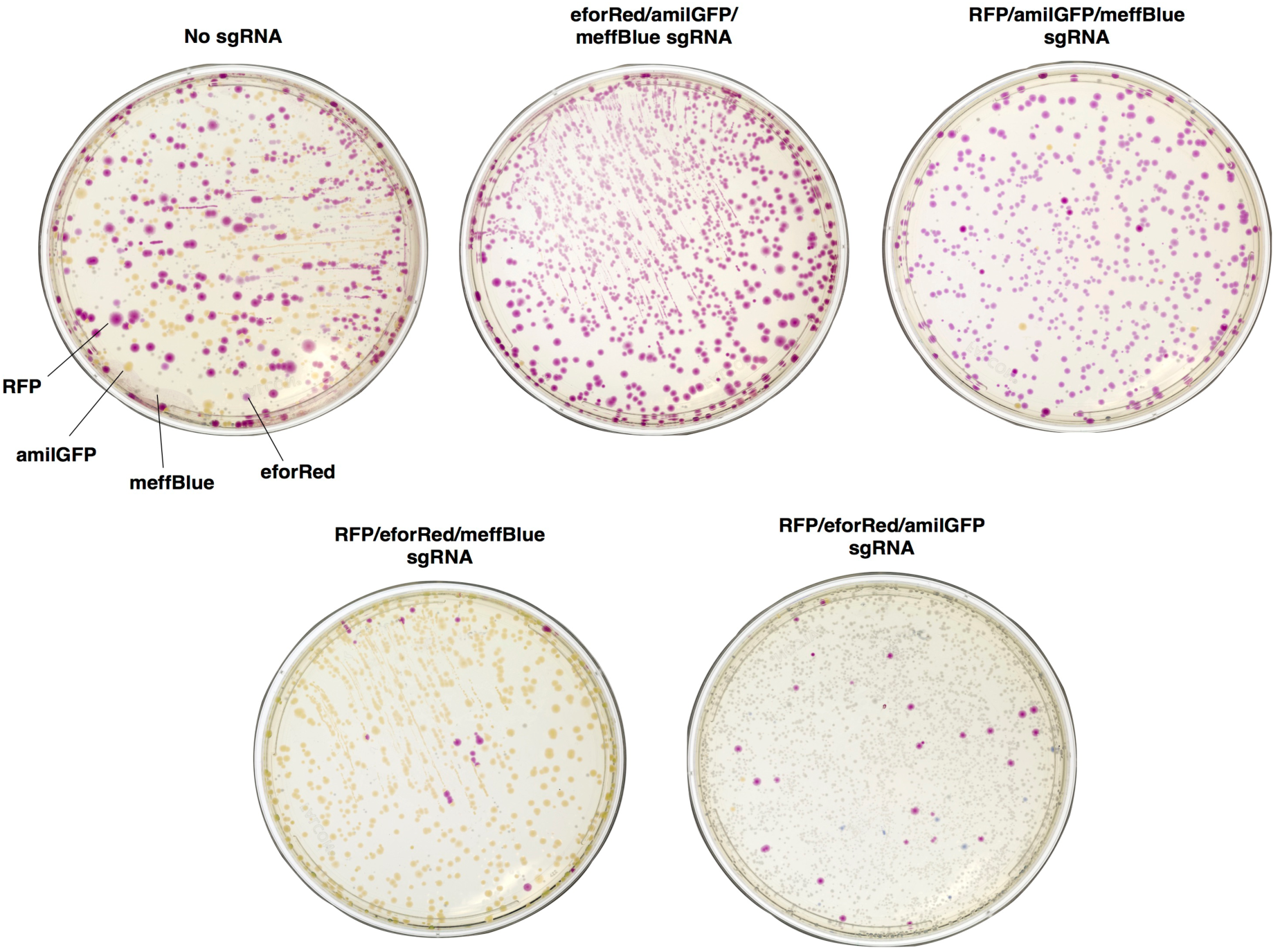
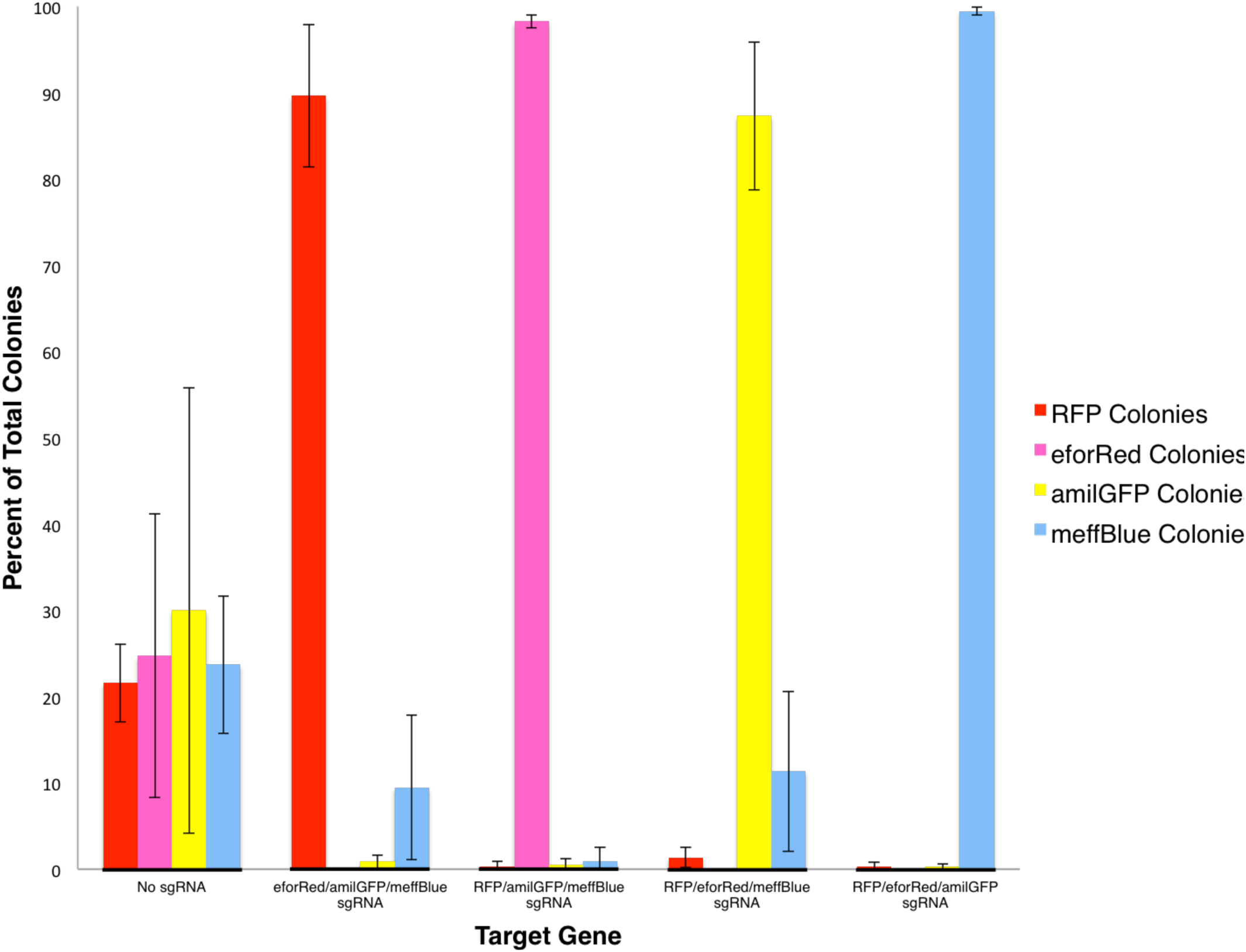
Cas9 cleavage of plasmids enhances transformation selectively. (A) Overview of CRATER method selecting for a chromogenic plasmid in a mixed pool. After ligation of chromogenic genes into the pSB1C3 backbone, plasmids are combined with recombinant *S pyogenes* Cas9 and sgRNAs specific to unwanted products. These products are cleaved *in vitro* into linear DNA, and are transformed along with the uncleaved plasmid. Intracellular exonucleases further cleave the linear DNA, leaving only the desired plasmid (in this case *amilGFP*) intact. (B) Plates of *E. coli* expressing color-producing proteins 48hr after transformation with a mixture of four plasmids. Before transformation, the plasmid mix was digested with Cas9 and either a mixture of three sgRNAs, or with a sgRNA-free control (No sgRNA). (C) Percentage of total colonies on plates with each color. Absent bars represent 0 colonies of that color found. Error bars represent SD, n = 3. Percentage values are shown in Supplemental Table S3.

**Figure 2.**
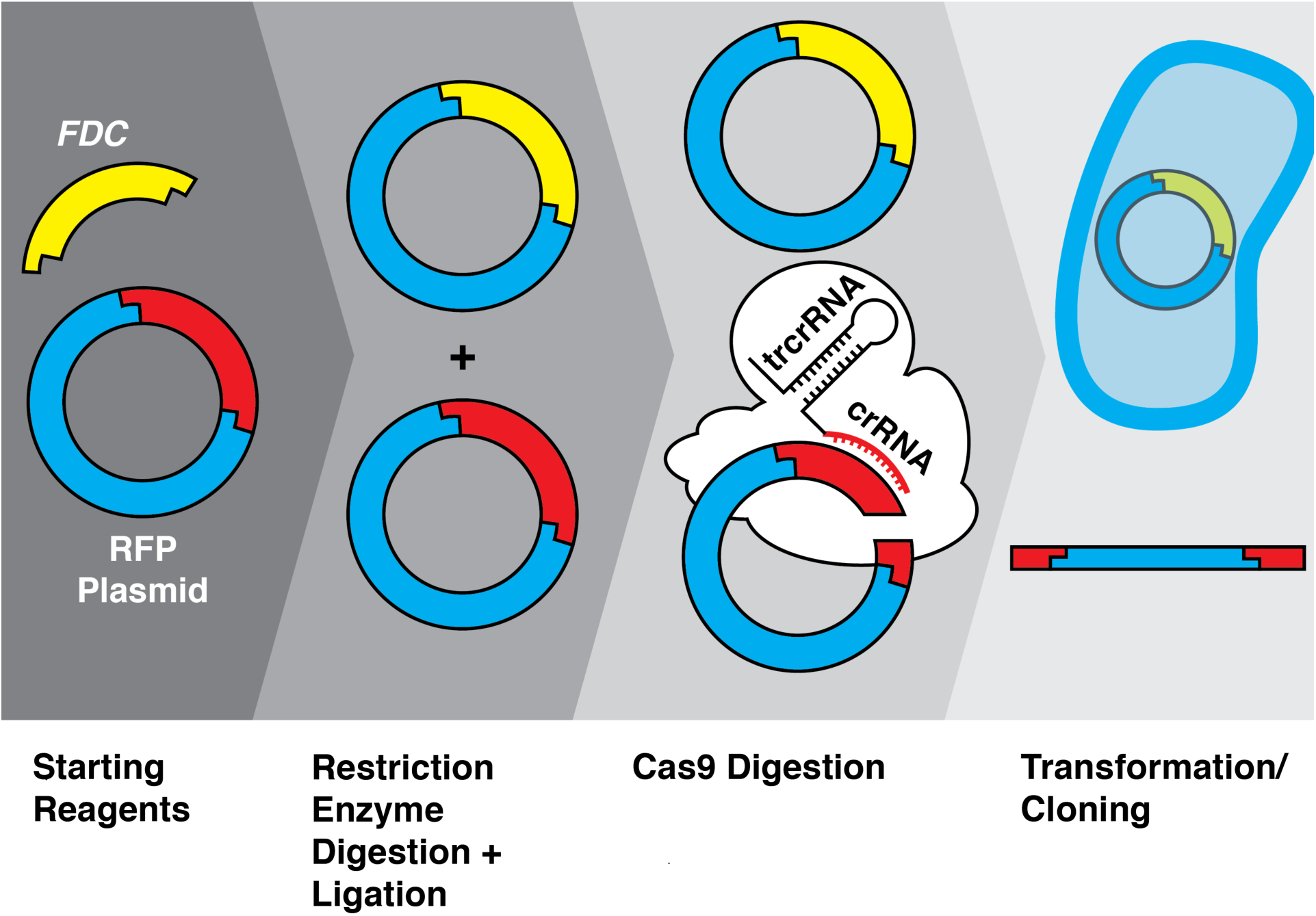
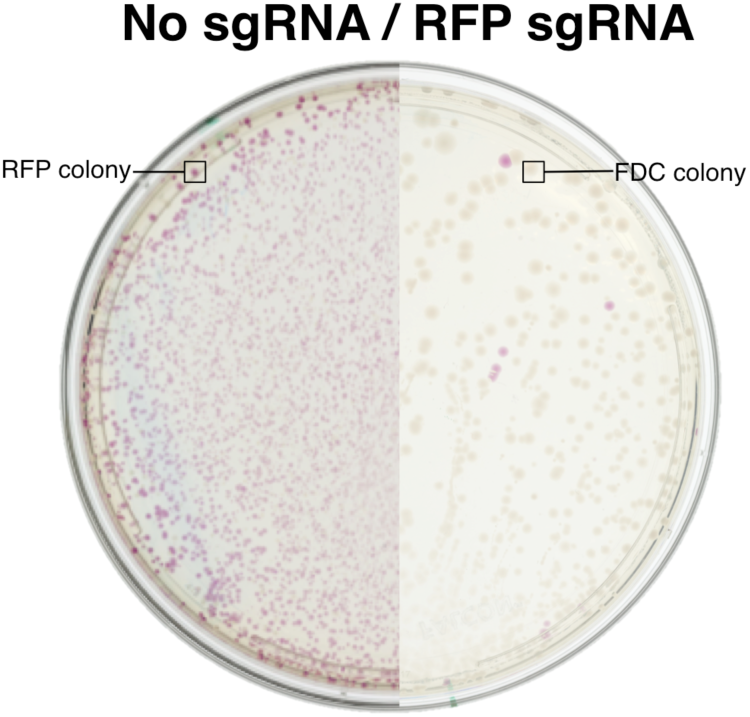
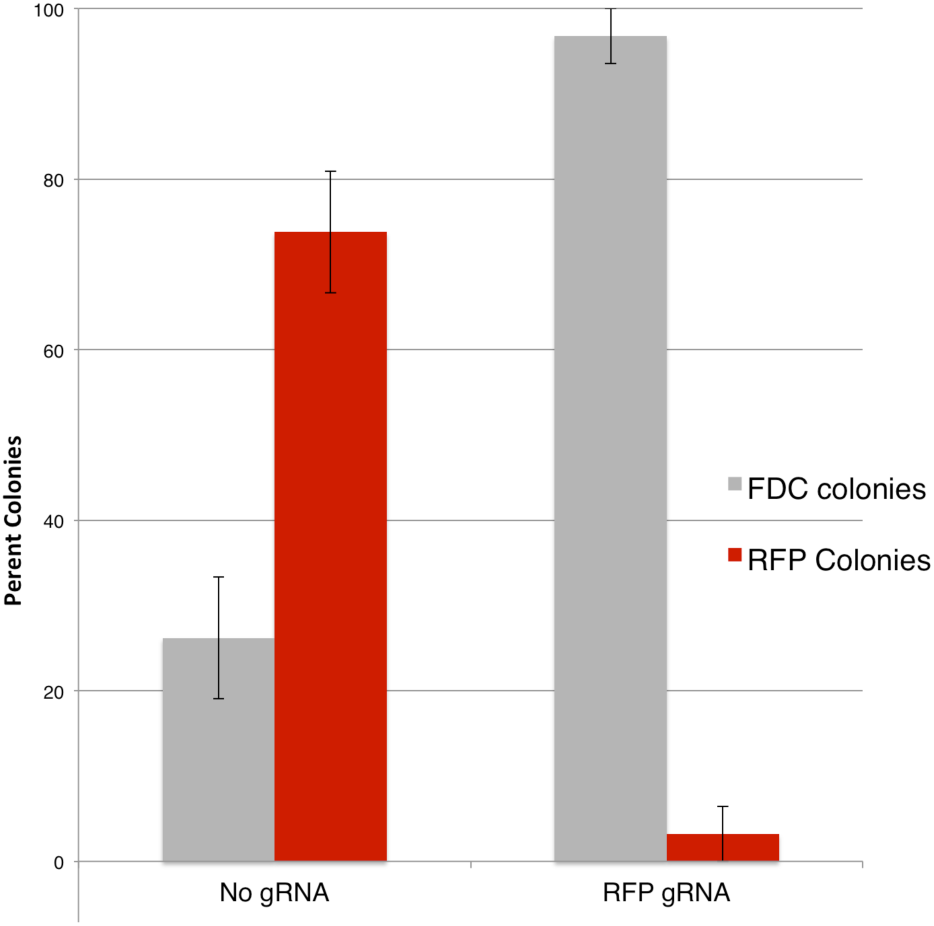
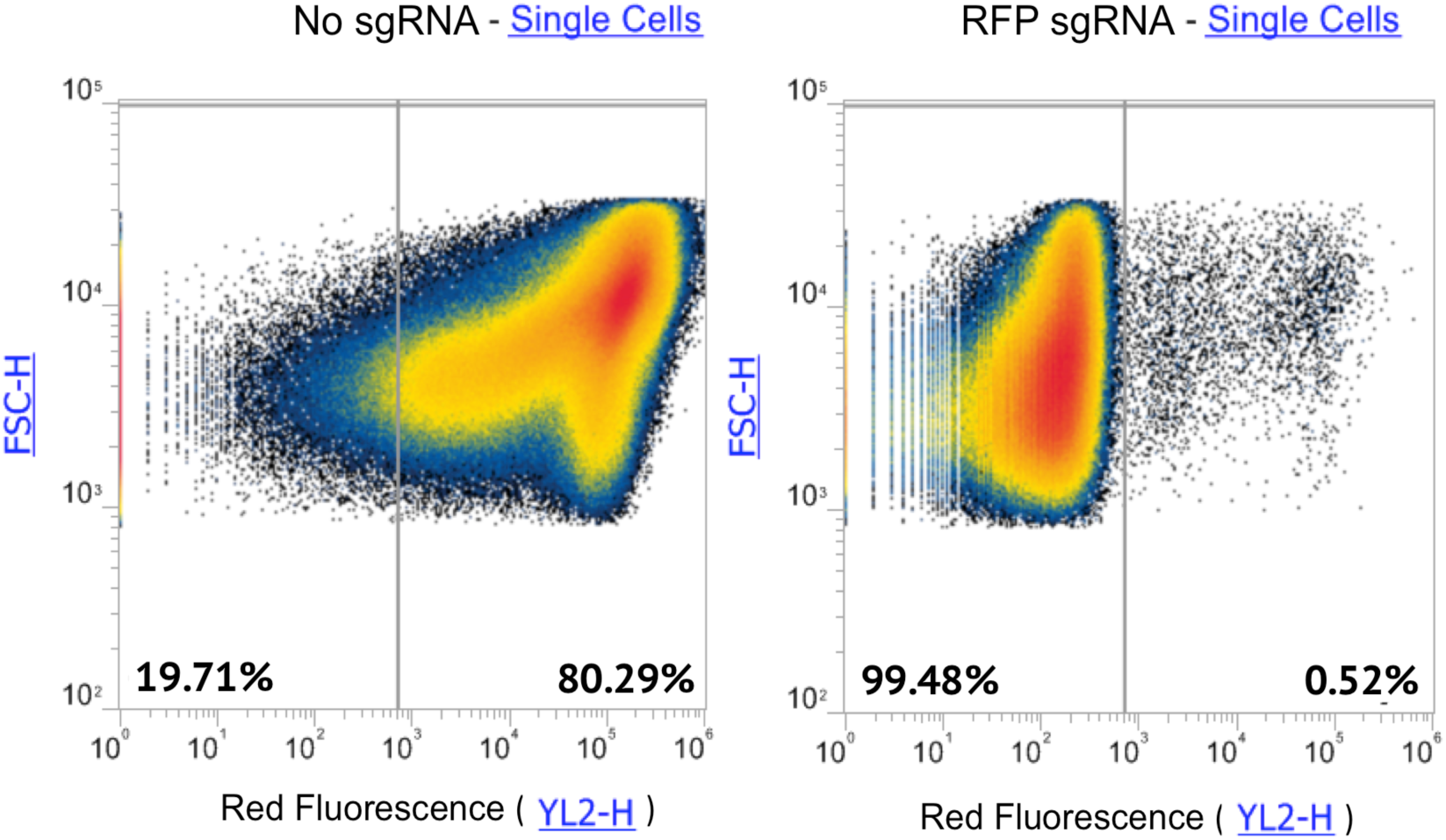
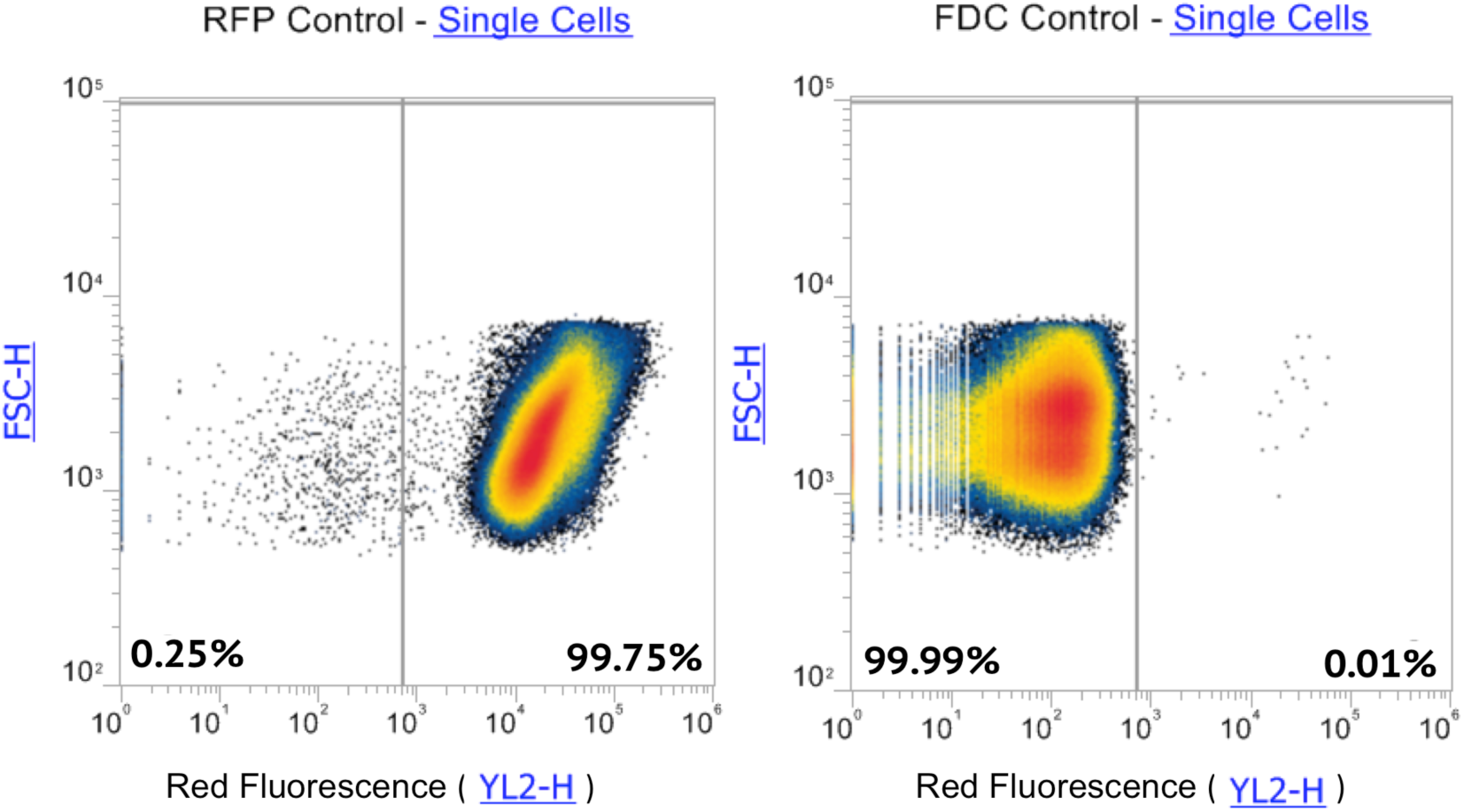
CRATER can selectively prevent the transformation of unwanted ligation products. (A) Overview of CRATER method selecting for FDC plasmid. (B) *E. coli* transformed with FDC and RFP mixed-ligation products, after either Cas9 digestion targeting the RFP gene or with a sgRNA-free control (No sgRNA). White colonies represent successful ligation of the FDC gene, while red colonies represent the unwanted RFP re-ligation product. (C) Percentage of total colonies on the plates with each insert, n = 2. Error bars represent SEM, p < 0.0004 according to two-way ANOVA test.(D) Flow cytometry data of the same transformations used in A and B after growing 24hr in liquid culture. (E) Flow cytometry data for pure RFP and pure FDC culture used to set the red fluorescence gate for C. The gates used to select for single cells are shown in Supplementary Figure S2.

### Cas9 cleavage can remove unwanted ligation products

We next sought to demonstrate that CRATER can remove unwanted plasmids created during ligation. As a basic example, we began with a plasmid containing an RFP gene and digested the gene out of its vector backbone with EcoRI and SpeI. We then added a sticky end compatible preparation of the Ferulic Acid Decarboxylase (FDC) gene, a gene that produces no color and represents the target for cloning, into the mix of EcoRI/SpeI digested RFP insert and vector backbone. Since the RFP insert was never removed from the restriction enzyme digestion, we predicted that a significant amount of the vector backbone would ligate with the original RFP insert instead of the FDC gene if CRATER were not utilized.

We transformed *E. coli* with the mixture of RFP and FDC plasmids subjected to CRATER using RFP-specific sgRNA or a sgRNA-free control. CRATER using RFP-specific sgRNA dramatically increased the percentage of colonies with the desired FDC insert when the transformed cells were plated (Fig. 3A, B). To corroborate this result, we analyzed the same transformed cells grown in liquid culture using flow cytometry. Applying CRATER reduced the percentage of red colonies from 80% to 0.5% in flow cytometry (Fig. 3C) supporting our solid culture results (Fig. 3B). Looking at the numbers of successful FDC transformations (Supplemental table 4), using CRATER increased our transformation efficiency from 20% to 97%.

### Discussion

By experimenting with chromogenic and fluorescent protein gene inserts, we have demonstrated the ability of Cas9 to digest and prevent the undesired transformation of plasmids in mixed-ligation pools. This technique is one of the many new applications of the recently discovered CRISPR/Cas system and can be used to augment existing methods for manipulating recombinant DNA.

CRATER will save time when manipulating DNA and constructing plasmids. Particularly in cases when transformation efficiency is very low, such as when transforming large genes (>10 kb) into plasmids, increasing efficiency will reduce the number of colony PCR amplifications and sequencing reactions required to find a colony with the desired insert. In addition to streamlining low-efficiency cloning experiments, CRATER could allow for tighter control of gene order and orientation during multi-component ligations. Designing sgRNAs that target reverse-oriented inserts could facilitate gene orientation control even when restriction enzymes give rise to complementary sticky ends. Moreover, sgRNA specificity could allow for this same level of control when inserting multiple genes into the a single vector. By designing sgRNAs that target unwanted orders and orientations of gene inserts, efficient construction of complex plasmids is made possible.

As Gibson assembly can also be used to efficiently construct plasmids with multiple genes, we wanted to discern whether the CRATER method was comparable to Gibson in terms of time and cost. To this end, we conducted a market study of biotechnology firms including New England Biolabs, Life Technologies, ElimBio, DNA2.0, and Integrated DNA Technologies (IDT). As of August 2015, the market price of a Gibson Assembly Kit was $630.00, with primers pairs ranging from $10 to $30 per pair. The combined market price of sgRNA synthesis materials and Cas9 digestion materials ranged from $708.80 to $874.30, assuming only one type of sgRNA was needed. Each additional sgRNA would cost $47.40. Both Gibson assembly and Cas9 digestion costs are for 50 reactions worth of materials, while the sgRNA synthesis cost is for 15-75 µg of sgRNA. Based on these estimates, CRATER appears to be roughly equivalent in cost to Gibson assembly, and is particularly useful for when fragment termini instability or repetitive DNA sequences prevent Gibson from being used, Gibson primer design results in primer dimer or hairpin formation, or when only one stock sgRNA is needed to prevent a particularly common unwanted ligation product. A good example of this latter case is plasmid backbones with standardized restriction sites, such as the PSB1C3 backbone used in the iGEM BioBrick Registry. This plasmid contains XbaI and SpeI restriction sites, so sgRNAs that target XbaI/SpeI-scarred re-ligated plasmids could be mass-produced and distributed with the standard backbone.

### Methods

#### Plasmids

All plasmids were obtained from the BioBrick Registry. BioBrick numbers, sizes, and descriptions are provided in Supplementary Table 1.

#### DNA Quantification and Sequencing

DNA concentrations were determined using the NanoDrop 2000 Spectrophotometer (Thermo Scientific, Waltham, MA, USA). Sequencing of DNA samples was completed by Elim Biopharmaceuticals, Inc. (Hayward, CA, USA).

#### sgRNA Preparation

The *t*tracrRNA reverse template primer, along with crRNA forward primers, were ordered from Elim Biopharmaceuticals, Inc. The 10 PCR primers used are shown in Supplementary Table 2. Single guide-RNA templates were PCR amplified from these primers in a 50 µL reaction, with initial denaturation at 98ºC for 30 seconds, annealing at 62ºC for 15 seconds, and elongation at 72ºC for 10 seconds, repeating for 10 cycles. The templates were isolated via the Epoch Life Sciences Inc. (Missouri City, TX, USA) PCR cleanup protocol. Transcription of sgRNAs was accomplished using the HiScribe™ T7 High Yield RNA Synthesis Kit (New England Biolabs, Ipswich, MA, USA). Last, sgRNAs were purified using the Life Technologies (Carlsbad, CA, USA) RNA extraction protocol.

#### Restriction Enzyme Digestion and Ligation

Single restriction enzyme digestion with PstI was accomplished using the New England Biolabs protocol. The MinElute Reaction Cleanup Kit (Qiagen, Limburg, Netherlands) was used to purify restriction enzyme digests. Restriction enzyme digests were ligated using the New England Biolabs T4 Ligation protocol.

#### CRISPR/Cas9 Digestion

The RFP and chromogenic plasmids were digested using the New England Biolabs *in vitro* Cas9 Digestion protocol, with the modification that 2 µL of 1µM Cas9 nuclease was added to the reaction instead of 1µL. When multiple sgRNAs were added to the same mixture, a 300 nM solution containing the sgRNAs was prepared in advance, and 3 µL of this solution was added to the reaction mixture.

#### Transformation

The PSB1C3 BioBrick plasmid backbone was used as a vector with chloramphenicol selection, and insert RFP and GFP genes were taken from the Biobrick Registry (BBa_J04450 and BBa_I13522, respectively). *E. coli* NEB5α chemically competent cells were purchased from New England Biolabs. Transformants were plated on LB plates with chloramphenicol selection by adding 50 µL of transformant mixture to the plate and spreading evenly using glass beads. 20 µL of transformants were also incubated in 3 mL of LB broth with chloramphenicol selection to analyze on the flow cytometer. Both plates and liquid cultures were grown at 37ºC for ~16 hours.

#### Plate Imaging

Plates were photographed using a Canon EOS 5D Mark II, Canon 100mm f/2.8 macro lens, and a fluorescent white light box.

#### Fluorescent Measurement

Liquid cultures of transformed *E. coli* were analyzed using Life Technologies Attune NxT Acoustic Flow Cytometer. 150 µL of each 5X dilute liquid culture was drawn at 12.5 µL/second until at least 500,000 events of single cells were collected (Supplementary Figure S2).

## Author Contributions

L.J.R. conceived the experiments. L.J.R., K.F., A.J.M., and J.D.S. oversaw the experiments and edited the manuscript. D.T.G., K.A.T., J.R.T., T.D.D., T.J., D.K., F.T., and D.X. performed the experiments and analyzed the data. D.T.G., D.K., K.A.T., J.R.T., C.C., E.L., and T.N. wrote the manuscript. All authors approved the final manuscript.

## References

Bhaya, D., Davison, M. & Barrangou, R. CRISPR-Cas systems in bacteria and archaea: versatile small RNAs for adaptive defense and regulation. Annu. Rev. Genet. 45, 273–297(2011).

Wiedenheft, B., Sternberg, S.H. & Doudna, J.A. RNA-guided genetic silencing systems in bacteria and archaea. Nature 482, 331–338 (2012).

Gasiunas, G., Barrangou, R., Horvath, P. & Siksnys, V. Cas9-crRNA ribonucleoprotein complex mediates specific DNA cleavage for adaptive immunity in bacteria. Proc. Natl. Acad. Sci. USA 109, E2579–E2586 (2012).

Jinek, M. et al. A programmable dual-RNA-guided DNA endonuclease in adaptive bacterial immunity. Science 337, 816–821 (2012).

F. J. Mojica, C. Díez-Villaseñor, J. García-Martínez, C. Almendros, Short motif sequences determine the targets of the prokaryotic CRISPR defence system. Microbiology 155, 733 (2009).

Conley E C, Saunders V, Saunders J R. Deletion and rearrangements of plasmid DNA during transformation of *Escherichia coli* with linear plasmid molecules. Nucleic Acids Res. 1986;14:8905–8917.

